# Ethoflow: computer vision and artificial intelligence-based software for automatic behavior analysis

**DOI:** 10.1101/2020.07.23.218255

**Authors:** Rodrigo Cupertino Bernardes, Maria Augusta Pereira Lima, Raul Narciso C. Guedes, Gustavo Ferreira Martins

**Author notes:** Corresponding author: R.C. Bernardes; E-mail addresses.

## Abstract

1. Manual monitoring of animal behavior is time-consuming and prone to bias. An alternative to such limitations is the use of computational resources in behavioral assessments, such as a tracking system, to facilitate accurate and long-term evaluations. There is a demand for robust software that addresses analysis in heterogeneous environments (such as in field conditions) and evaluates multiple individuals in groups while maintaining their identities.
2. The Ethoflow software was developed using computer vision and artificial intelligence (AI) tools to automatically monitor various behavioral parameters. A state-of-the-art object detection algorithm based on instance segmentation was implemented, allowing behavior monitoring in the field under heterogeneous environments. Moreover, a convolutional neural network was implemented to assess complex behaviors, thus expanding the possibilities of animal behavior analyses.
3. The heuristics used to automatically generate training data for the AI models are described, and the models trained with these datasets exhibited high accuracy in detecting individuals in heterogeneous environments and assessing complex behavior. Ethoflow was employed for kinematic assessments and to detect trophallaxis in social bees. The software runs on the Linux, Microsoft Windows, and IOS operating systems with an intuitive graphical interface.
4. In the Ethoflow algorithm, the processing with AI is separate from the other modules, which facilitates kinematic measurements on an ordinary computer and the assessment of complex behavior on machines with graphics processing units (GPUs). Thus, Ethoflow is a useful support tool for applications in biology and related fields.

## 1 Introduction

Behavioral studies are critical to understanding the fundamental aspects of animal ecology (Anderson & Perona, 2014; Dell et al., 2014). The assessment of animal behavior by visual inspection is limited and subjective, and does not allow observations over long periods (Noldus, Spink, & Tegelenbosch, 2002). The use of computational tools in behavioral assessments, such as automatic tracking systems, allows accurate and long-term evaluations of animals (Dell et al., 2014; Valletta, Torney, Kings, Thornton, & Madden, 2017). Calculation of important kinematic measurements, including the tracked distance, is feasible when tracking animals, and the evaluation of complex behaviors can provide relevant insights about animal biology. For example, the evaluation of complex behaviors among social insects, such as changes in grooming and trophallaxis, is important for understanding their response to stress agents such as pesticides (Gandra, Amaral, Couceiro, Della Lucia, & Guedes, 2016; Boff, Friedel, Mussury, Lenis, & Raizer, 2018).

Robust systems are needed for animal monitoring in heterogeneous environments, such as in the field (Dell et al., 2014). The greatest challenge in heterogeneous environments involves the extraction of target objects from the background (segmentation) (Zou, Shi, Guo, & Ye, 2019). Animal tracking software operates by background subtraction or thresholding (Yamanaka & Takeuchi, 2018; Sridhar, Roche, & Gingins, 2019). As these approaches require video recordings with sufficient contrast between the object and homogeneous background, they are not applicable in the field. Software using artificial intelligence (AI) can be sufficiently robust for assessments in heterogeneous environments, as AI models can be trained to learn the detection of target objects in different environments (He, Gkioxari, Dollár, & Girshick, 2018).

Convolutional neural networks (CNNs) are the most conventional AI models employed in computer vision tasks. While these models have exhibited outstanding performance in computer vision applications and tracking software, including idtracker.ai and DeepLabCut, which use AI models in their algorithms, they still exhibit some limitations. The idtracker.ai software applies a CNN to maintain the identity of individuals in a group, but its application is limited to homogeneous environments (Romero-Ferrero, Bergomi, Hinz, Heras, & de Polavieja, 2019). While DeepLabCut uses a CNN for animal pose estimation and tracks parts of objects in heterogeneous environments (Nath et al., 2019), it does not maintain the identity of individuals in a group.

Given the potential applications of AI and the demand for a robust system that fulfills the requirement for studying animal behavior (Dell et al., 2014), the Ethoflow software was developed. AI was incorporated into Ethoflow for object detection, enabling evaluations of the complex behaviors of individuals in groups living in heterogeneous environments. To validate Ethoflow, bioassays with two species of eusocial bees were performed. In addition, parameters associated with the performance of the Ethoflow software were evaluated during its execution.

## 2 Software features

The Ethoflow software runs on Linux, Microsoft Windows, and IOS operating systems. This software is registered with the Brazilian National Institute of Intelectual Property (Instituto Nacional de Propriedade Industrial, INPI, Ministério da Economia, Brazil, reg. no. BR 51 2020 000737-6). Ethoflow is user-friendly and does not require the use of line commands because of the intuitive graphical user interface (GUI) (Fig. 1A). There are three tabs (Settings, Analyses, and Deep analyses) on the Ethoflow GUI. Under the ‘ Settings’ tab, the parameters can be set. Once the parameters have been set, they can be saved as a protocol (as a .txt file) and loaded for the next step. When setting the parameters, the interface allows real-time monitoring of the effects of parameter changes; for example, detected objects (i.e., animals) will be marked with a red mask. A region of interest (Fig. 1A; blue box on the bottom right) can also be defined to assess how long the individuals stayed in this region. The ‘Analyses’ tab corresponds to the video processing step and renders images to train the instance segmentation model**Erro! Fonte de referência não encontrada.**. During video processing, the user can visually monitor the processing in real time. Following completion of processing, the program prints the processing speed and detection rate on the GUI. Under the ‘Deep analyses’ tab, models for complex behavior analysis are loaded, which allows the recognition of several specific behaviors on the condition that an AI model is set up. The median and standard deviation of some measurements of the body of individuals (for e.g., area and length) can also be calculated. These parameters can serve as a basis for generating labeled images to train the specific behavior model and assist in the protocol definition.

**Figure 1.**
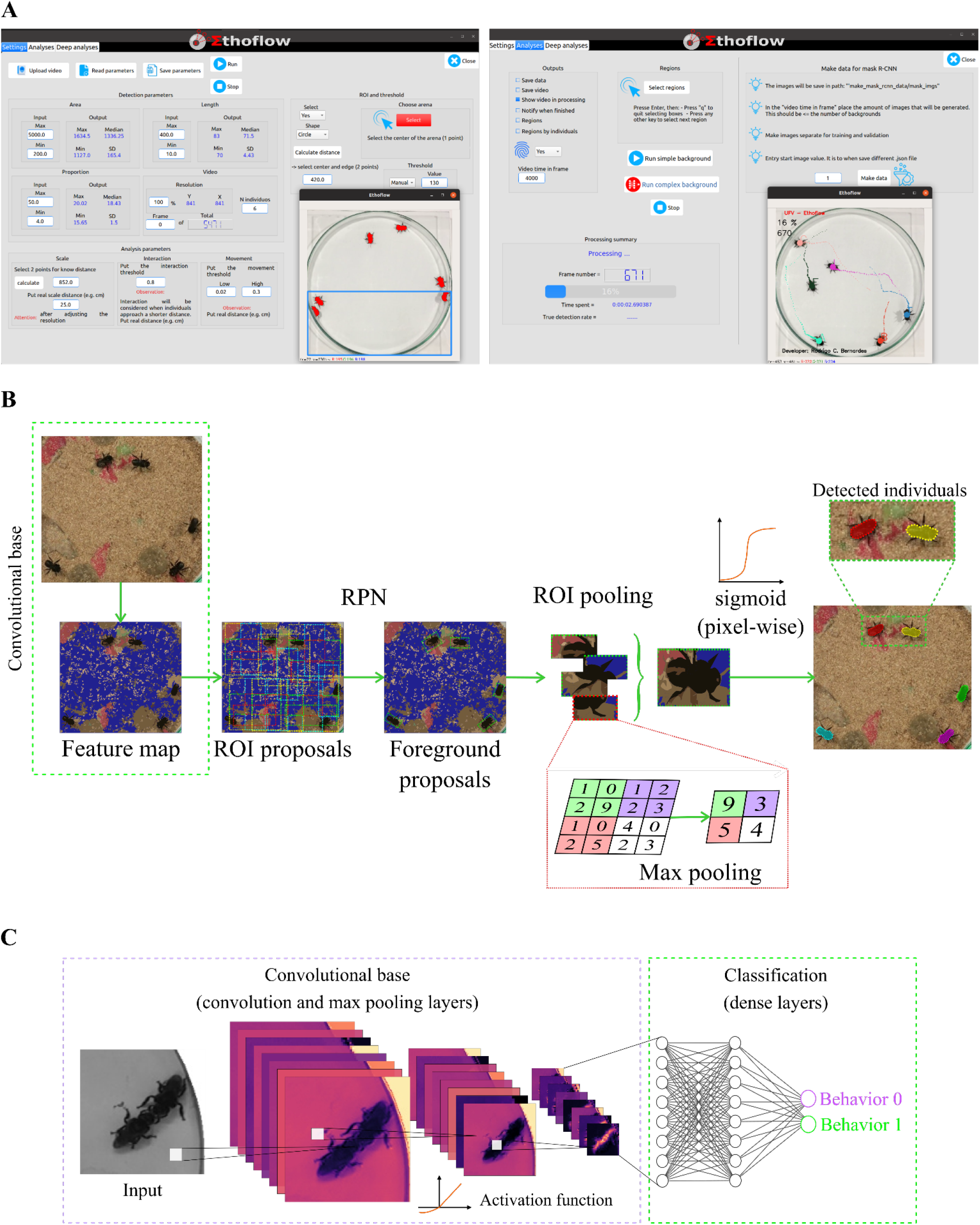
Features of the Ethoflow software. (A) Graphical user interface (GUI). In the left panel (Settings tab), the detected individuals are marked with a red mask after adjusting the parameters (protocol definition). Blue box on the bottom right shows the selected region of interest, which is used to assess how long the individuals remain inside the defined region. In the right panel (Analyses tab), monitoring of the processing step is demonstrated. Each individual is randomly assigned an identity and color. In this example, the analyzed objects correspond to workers of the stingless bee species *Melipona quadrifasciata* present in a Petri dish (15 cm diameter, 2 cm height). (B) Diagram depicting the operations involved in instance segmentation. The input passes through a convolutional base for feature extraction, leading to the generation of a feature map. The region proposal network (RPN) is applied, which provides several candidate boxes (ROI proposals). As several ROIs are generated, the model classifies these boxes into foreground proposals (objects) and backgrounds. ROI pooling is applied to standardize the size of the foreground proposals, slicing each foreground into a fixed amount of parts, and max pooling is applied to standardize the size. Then, the boxes labeled as real objects (the individuals) are instantiated using a pixel-wise sigmoid function. (C) Workflow of the convolutional neural networks used in Ethoflow to assess complex behaviors. In the convolutional base, the input passes through the convolutional and max pooling layers for feature extraction. Then, behavior classification is performed in the dense layers. The activation function is applied to the output of each layer to introduce non-linearity.

In the Ethoflow algorithm, multi-threaded reading was implemented. This avoids the delay between frame reading and other processing steps of the algorithm, whereby frames are always available, thus making it possible to obtain high rates in frames per second (fps) (Supporting Information A. 1). In addition, the AI processing is separate from the other modules in the Ethoflow algorithm. Therefore, Ethoflow enables kinematic measurements on an ordinary computer and the assessment of more complex behavior using a GPU.

Manual and automatic thresholds were applied in Ethoflow to detect individuals. In manual thresholding, the threshold value is defined by the user. One of the automatic thresholding options is based on Otsu’s method (Otsu, 1979), wherein the optimal threshold minimizes the within-class variance (Supporting Information A.3). The other automatic thresholding option involves instance segmentation (IS), which is based on the Mask R-CNN AI model (He et al., 2018). This state-of-the-art model for object detection allows Ethoflow to work in field conditions, detecting individuals in a heterogeneous background (Fig 1. B). The hyperparameters and architecture of the IS model are based on the implementation reported by Abdulla (2017) (Supporting Information A.3). The IS model should be trained to learn to detect the animal of interest. Thus, a heuristic was used to automatically generate the training data. This heuristic functions by extracting the contours of individuals in a homogeneous background and subsequently pasting them in a heterogeneous background. (Supporting Information A.3).

Ethoflow monitors animals in groups, maintaining the identity of individuals. In this step of the algorithm, the nonhierarchical clustering k-means is applied to separate merged individuals. Subsequently, a combinatorial optimization algorithm is applied, which provides the optimal identity assignment between individuals (Supporting Information A.5). Ethoflow records the coordinates of movement in time and calculates various kinematic parameters associated with the behavior of individuals and groups (Supporting Information A.6). Moreover, Ethoflow measures specific behaviors using a CNN model (Fig. 1C); different hyperparameter configurations were tested to define the CNN model, which can be used to recognize binary behaviors (Supporting Information A.7). The data used to train the CNN model was also generated with a heuristic based on the animal body size and length (Supporting Information A.7).

## 3 Performance and applications

Ethoflow was trained to detect the bee *Melipona quadrifasciata* in several field conditions (Supporting Information B.1). The proposed model was efficient in detecting all animals in heterogeneous backgrounds with high precision (average precision ± standard error = 0.916 ± 0.02; Fig. 2A). Ethoflow was also trained to learn the detection of trophallaxis, the complex social behavior of food exchange among nestmates, in *M. quadrifasciata* (Supporting Information B.1). This model exhibited high accuracy in the validation process (global accuracy = 92.13%, Kappa index = 0.84, Z = 24.74, Fig. 2B).

**Figure 2.**
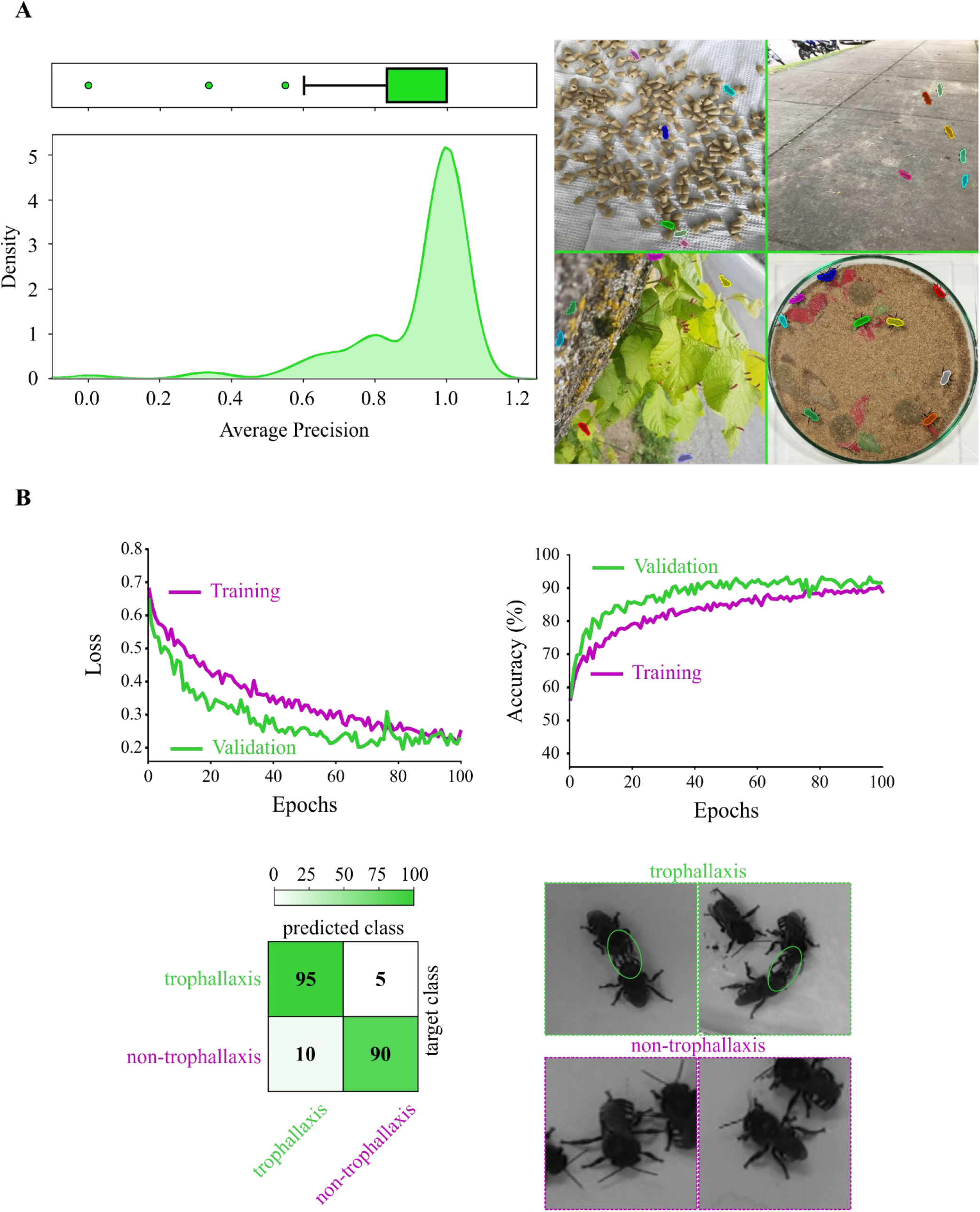
Performance of the AI models used in Ethoflow. (A) Object detection in heterogeneous backgrounds based on instance segmentation (IS). The high average precision (left panel; *n* = 100) implies that the model precisely detects real objects in the scenes with no false positives, as demonstrated by (right panel) the detected objects (*Melipona quadrifasciata* bees) with masks in random colors. (B) The training process of the CNN model (top panel) and cross-validation (confusion matrix; bottom left panel) (*n* = 127) for the monitoring of trophallaxis (green circles) in bees.

To validate Ethoflow, a behavioral assay was performed with the stingless bee species *M. quadrifasciata* and *Partamona helleri* (Supporting Information B.2) and different kinematic variables were measured (definitions of the variables are detailed in Supporting Information A.6). In both species, the centrality increased with the polarization of the group (F _1, 35_ = 25.1, *p* < 0.0001) and decreased with milling (F _1, 35_ = 46.2, *p* < 0.0001) (Fig. 3A). Meandering was influenced by the statistical interaction between the variables resting and bee species (F _1, 33_ = 4.71, *p* = 0.037; Fig. 3B). Moreover, a difference between species was observed in the tracked distance (F _1, 35_ = 13.6, *p* = 0.0008; Fig. 3C).

**Figure 3.**
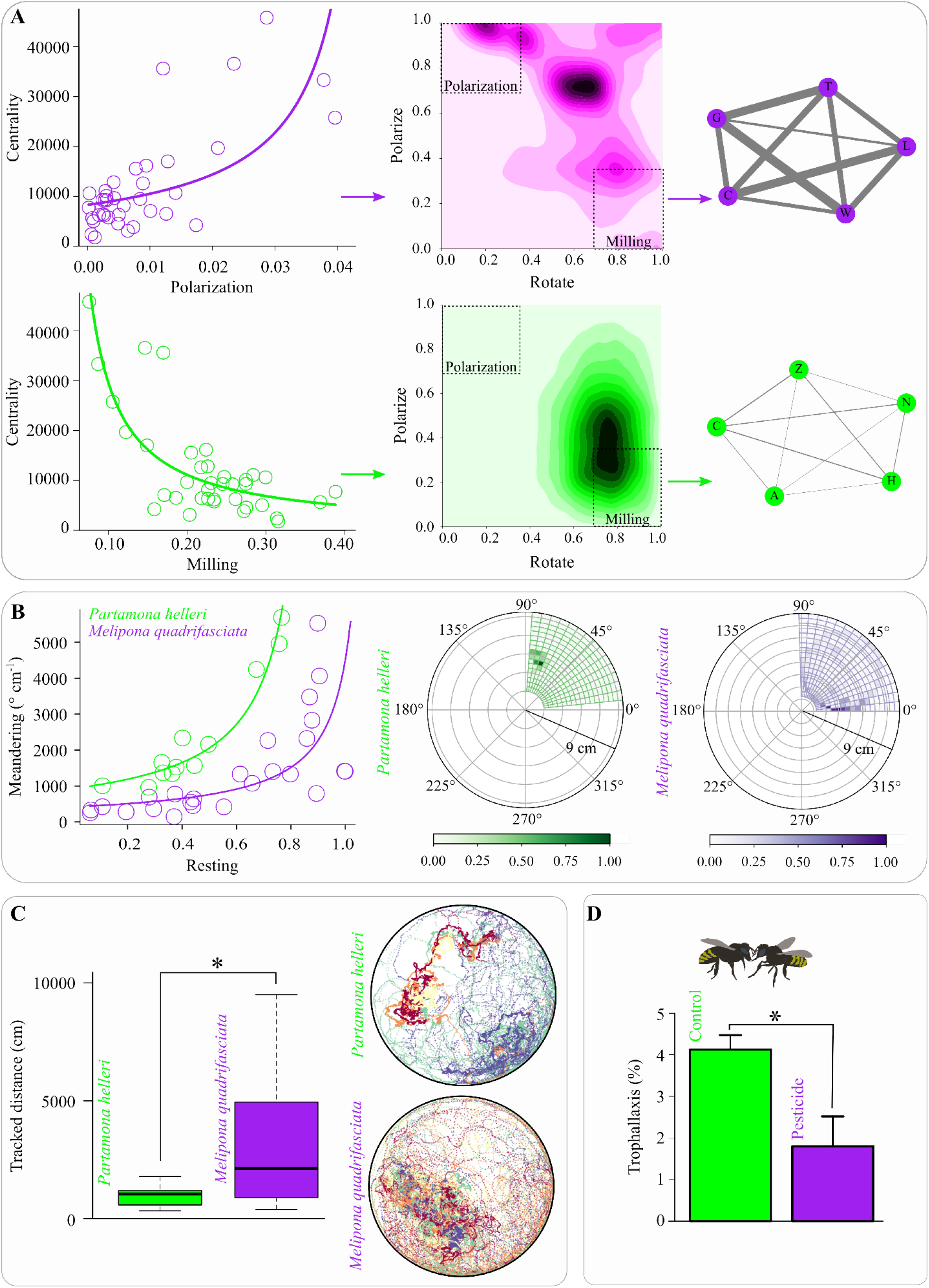
Behavioral assessments conducted using the Ethoflow software. (A) Association between centrality and group dynamics polarization (top panel) and milling (bottom panel) (*n* = 37). The 2D density plots and network diagrams showed that in more polarized bee groups, a higher interaction exists among individuals, while this interaction is reduced in the milling groups. In the networks, the circles represent individuals and connections correspond to the edges, where their widths are proportional to the frequency of interactions. (B) Meandering behavior is associated with resting proportions (left panel) (*n* = 37) and histograms of polar coordinates (rays and azimuth angles) for the two bee species (right panel). (C) The tracked distance of the assessed bee species (*n* = 37). In group representative tracks, the track color reflects the individual identity (right panel). (D) Trophallaxis alteration in *M. quadrifasciata* after pesticide exposure (*n* = 60). * *p* < 0.05 in the generalized linear model. When an explanatory variable had no significant effect, the model was simplified, and the results were plotted as a function of the significant variable.

A toxicological bioassay was performed with *M. quadrifasciata* to demonstrate the recognition of trophallaxis under pesticide stress conditions. The pesticide imidacloprid, which is usually associated with the decline of bees, was used (Lima, Martins, Oliveira, & Guedes, 2016). The exposure protocol was based on that reported by Botina et al. (2020) (Supporting Information B.2). Bees exposed to the pesticide exhibited significantly reduced trophallaxis (χ^2^ = 94.9, df = 58, *p* < 0.0001; Fig. 3D).

Considering the parameters associated with the software performance (Supporting Information B.3), Ethoflow achieves a high processing speed, reaching 300 fps (Fig. 4). In the homogeneous backgrounds, statistical interaction was observed between the variables video resolution and group size in the range of fps (F _1, 130_ = 12.81, *p* = 0.0005, Fig. 4A). The heterogeneous environment quantification was not influenced by the resolution or number of individuals (F _1, 28_ = 0.81, *p* = 0.37, Fig. 4B). The fps decreased with an increase in the centrality of individuals (F _1, 38_ = 81.24, *p* < 0.0001, Fig. 4C). There was no significant effect on the number of individuals (F _1, 37_ = 0.009, *p* = 0.93), and no interaction was observed between the centrality and individuals (F _1, 36_ = 1.62, *p* = 0.21). In addition, the software exhibited high detection rates (Fig. 4D). A significant interaction was observed between the number of individuals and type of background (F _1, 94_ = 137.85, *p* < 0.0001), where an increase in the number of individuals had a greater influence on the heterogeneous environments. A few videos processed with Ethoflow are available in B.4 of the Supporting Information.

**Figure 4.**
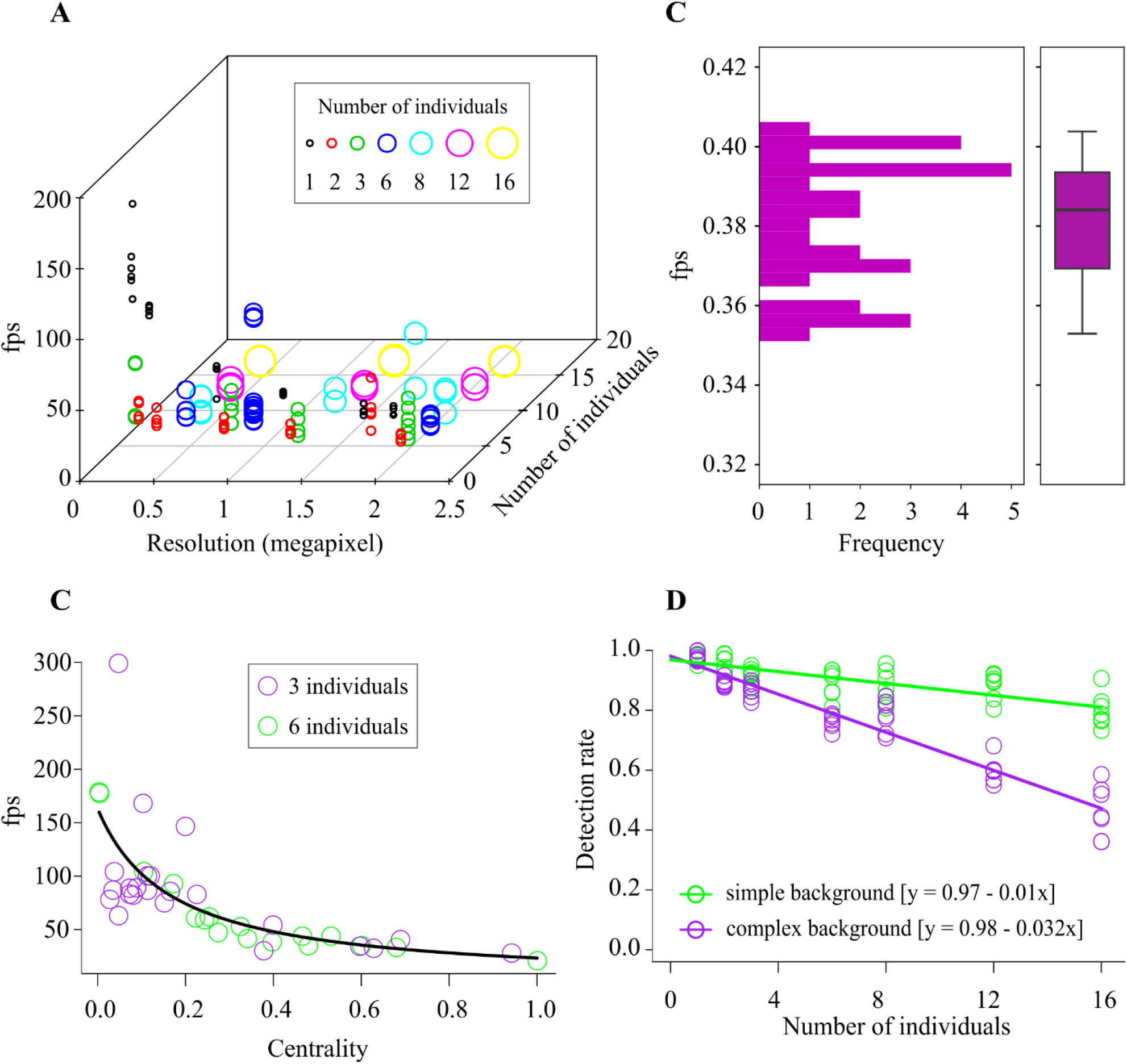
Quantification of the performance of Ethoflow. (A) Fps in response to the video resolution (in pixels) and number of individuals in homogeneous backgrounds; the dots (*n* = 134) represent the raw data. (B) Histogram of the fps in heterogeneous environments (*n* = 30). The box plot indicates the median and range of dispersion (lower and upper quartiles and outliers). (C) Fps in response to centrality. The proportion of group interaction per frame was used to quantify the centrality (*n* = 40). (D) Accuracy of the software as a function of the interaction between the number of individuals and type of environment (homogeneous and heterogeneous); the symbols represent the raw data (circles; *n* = 98). When an explanatory variable had no significant effect, the model was simplified, and the results were plotted as a function of the significant variable.

## 4 Conclusion

The Ethoflow software was developed using modern computer vision techniques and AI. This software exhibited consistent speed rates and processing accuracy. The developed software is suitable for behavioral assessments in heterogeneous environments, to track groups with individuals maintaining their identities, and can be trained to learn specific behaviors. Ethoflow was applied to biological assessments, and some possibilities of data analysis and representation were demonstrated with Ethoflow’s output. Accurate AI models have been implemented to expand the possibilities of animal behavior analyses to other fields, including the behavioral monitoring of domestic animals in precision livestock farming. Therefore, Ethoflow is a useful support tool for technical and scientific applications in biology and related fields.

## Acknowledgements

We thank the National Council of Scientific and Technological Development (CNPq 142206/2017-2 and 301725/2019-5) and CAPES-PROEX for financial support (Financial Code 001).

## Authors’ contributions

RCB developed the software, performed the experiments, analyzed the data; RCB and GFM wrote the manuscript. All authors designed the work and contributed to the drafts and gave final approval for publication.

## Data archiving statement

The data have been archived at https://github.com/bernardesrodrigoc/Ethoflow; DOI: https://doi.org/10.5281/zenodo.3956831

