## Supporting Information for "Ethoflow: computer vision and artificial intelligence-based software for automatic behavior analysis"

#### A. Software features (section A)

The Ethoflow software was developed using the Python language (version 3.7.3), including the image library OpenCV and the framework TensorFlow with Keras for AI models. Other libraries such as SciPy, Numpy, Pandas, and SciKit Learn were also used. The steps of the developed algorithm are shown in Fig. S1. The following subsections will provide further details on the different algorithm steps.

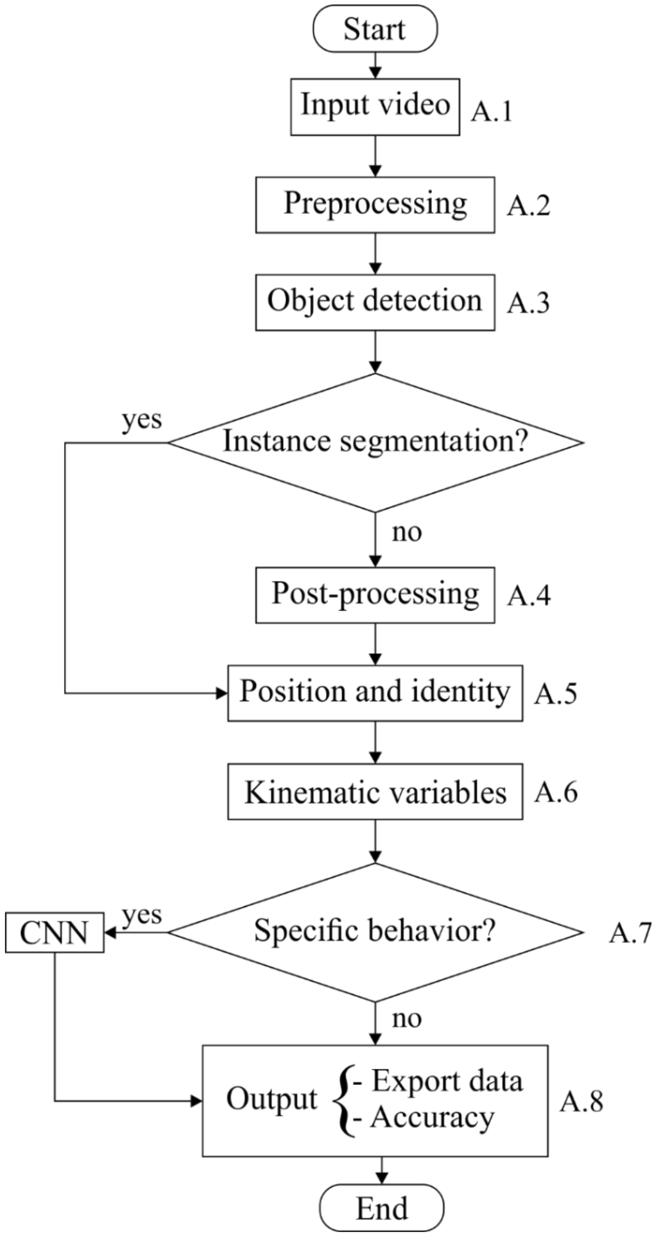


Figure S1. Block diagram of the Ethoflow algorithm. CNN represents a convolutional neural network.

#### A.1 Input video

The video is read by a thread that is independent of the processing thread, and the frames are stored in a stack (queue). This queue is a linear data structure that stores items in a "FIFO" (First In, First Out) manner (Fig. S2). Frames are exchanged between the reading and processing threads. This increases the processing speed, as frames are always present in the queue and ready for processing, and no time is spent waiting for the next frame to be read.

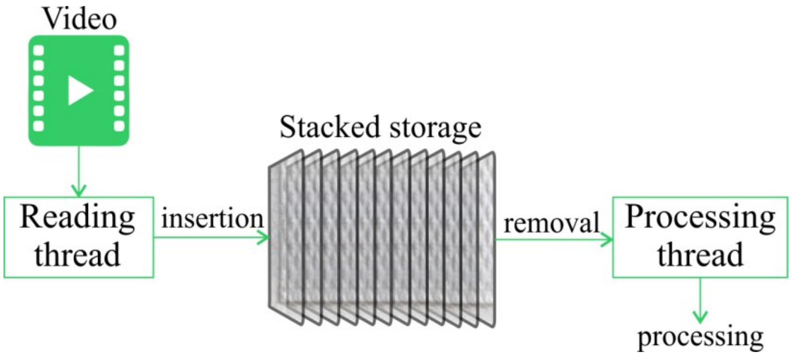


Figure S2. Representation of the processing steps in the multithreading feature of Ethoflow.

#### A.2 Preprocessing

If a region of interest is defined, the video will be processed to eliminate the regions that are not of interest to the user, which could compromise the analysis. The video is transformed into a virtual primary color system (color space XYZ). In this color space, the chromaticity (XZ) and luminance (Y) are coded separately. This results in a more uniform response to the variation in the luminosity. Then, grayscale transformation and normalization are applied to increase homogeneity between the frames. Smoothing (to eliminate noise) is also applied through a transformation in the spatial domain via convolution; an implementation based on the neighborhood median was employed.

#### A.3 Object detection

After preprocessing, thresholding is applied to segment the individuals of interest; both manual and automatic thresholds can be implemented. In manual thresholding, the classification of pixel $(x, y)$ is performed according to a global threshold defined by the user ($k$):

$$f(x,y) = \begin{Bmatrix} 1\ if(x,y) > k \\ 0\ if(x,y) \leq k \end{Bmatrix}. \quad (1)$$

One of the automatic thresholding options is Otsu's method (Otsu, 1979). This algorithm attempts to find a threshold value ($k$) that minimizes the within-class variances $c_0$ and $c_1$ (background and objects, respectively). If the set of gray levels of an image $L = \{1, 2, \cdots, l\}$ and the total number of pixels $N = \{n_1, n_2, \cdots, n_l\}$, then the probability of occurrence of a gray level $(p_i)$ is given by

$$p_i = \frac{n_l}{N}. \quad (2)$$

As the method is based on the normalized histogram,

$$\sum_{i=1}^{L} p_i = 1. \tag{3}$$

Thus, the probability of occurrence $(\omega_i)$, means $(\mu_i)$, and variances $(\sigma_i)$ of each class, are given by

$$\omega_0 = \sum_{i=1}^{k} p_i \text{ and } \omega_1 = \sum_{i=k+1}^{L} p_i, \tag{4}$$

$$\mu_0 = \frac{\sum_{i=1}^{k} i * p_i}{\omega_0} \text{ and } \mu_1 = \frac{\sum_{i=k+1}^{L} i * p_i}{\omega_1}, \tag{5}$$

$$\sigma_0^2 = \frac{\sum_{i=1}^{k} (i-\mu_0)^2 p_i}{\omega_0} \text{ and } \sigma_1^2 = \frac{\sum_{i=k+1}^{L} (i-\mu_1)^2 p_i}{\omega_1}. \tag{6}$$

The within-class $(\sigma_w)$ and between-class $(\sigma_b)$ variances are

$$\sigma_w^2 = \omega_0 \sigma_0^2 + \omega_1 \sigma_1^2, \tag{7}$$

$$\sigma_b^2 = \omega_0 \omega_1 (\mu_1 - \mu_0)^2. \tag{8}$$

The total variance is $\sigma_t^2 = \sigma_w^2 + \sigma_b^2$, and calculating the between-class variance improves the computational time because the variance between classes is based on first-order statistics (class means) (Otsu, 1979).

Instance segmentation (IS) is another type of automatic segmentation available in Ethoflow. ResNet-101 was the convolutional base used in this model (He, Zhang, Ren, & Sun, 2016), following the Mask R-CNN implementation (Abdulla, 2017). The detection confidence was set to 85%. This rate determines the threshold for which the foregrounds are passed to the classification and the mask is determined. Here, this lower threshold was interesting, as it is better to detect false objects compared with the failure to detect the true objects. False objects can be eliminated after morphological operations based on the user inputs.

The evaluation of the IS model was based on the average precision (*AP*) (Everingham, Van Gool, Williams, Winn, & Zisserman, 2010). To obtain this measure, the Intersection over Union (*IoU*) of the predicted bounding boxes (i.e., the *x, y* coordinates in the upper-left corner and width and height of the rectangular box around the object of interest) and target bounding boxes is calculated. Based on the *IoU*, the precision (Eq. 9) and recall (Eq. 10) can be calculated, using the true positives (*TP*), false positives (*FP*), and false negatives (*FN*) for the detected objects (*DO*) in a determined threshold (*x*) (Eq. 11).

$$\text{precision} = \frac{TP}{TP + FP} \tag{9}$$

$$\text{recall} = \frac{TP}{TP+FN}. \tag{10}$$

$$\begin{Bmatrix} \text{if } IoU \geq x, DO = TP \\ \text{if } IoU < x, DO = FP \\ \text{if the model fails to detect a target object}, DO = FN \end{Bmatrix} \tag{11}$$

There is a tradeoff between the precision and recall, wherein the higher the recall, the more the model tends to find all the target objects, i.e., a low *FN* value. However, an increase in the recall tends to decrease the precision, as it increases *FP*.

Considering equally spaced recall levels $n = (0, 0.1, \cdots, 1.0)$, interpolation is performed using the highest precision value for a given recall. Then, the *AP* is obtained from the interpolated values of the precision $\left(P_{interp}(r)\right)$:

$$AP = \frac{1}{n}\sum_{r=0}^{n} P_{interp}\,(r_i). \tag{12}$$

The inputs with bounding box positions, classes, and masks (pixel-wise positions of the objects) are required to train the IS model (He, Gkioxari, Dollár, & Girshick, 2018). The manual generation of these inputs is a laborious task. Then, a heuristic is developed to automatically generate these inputs based on several random backgrounds and a video in homogeneous conditions to detect objects using manual segmentation or Otsu's method. Frames are randomly sampled in the video and pass through the preprocessing and object detection stages of the algorithm (Fig. S3A). Then, the detected objects are "copied" and the contours are "pasted" into random backgrounds (Fig. S3B). Concomitantly, the bounding box, class, and mask of each object are saved in a dictionary with the following structure: Dictionary {image$_i$: {object$_j$: {box: {center: {x,y}, width, height}; class:{target}; mask:{all points (x,y)}}}}. The number of images that is generated is configured by the user.

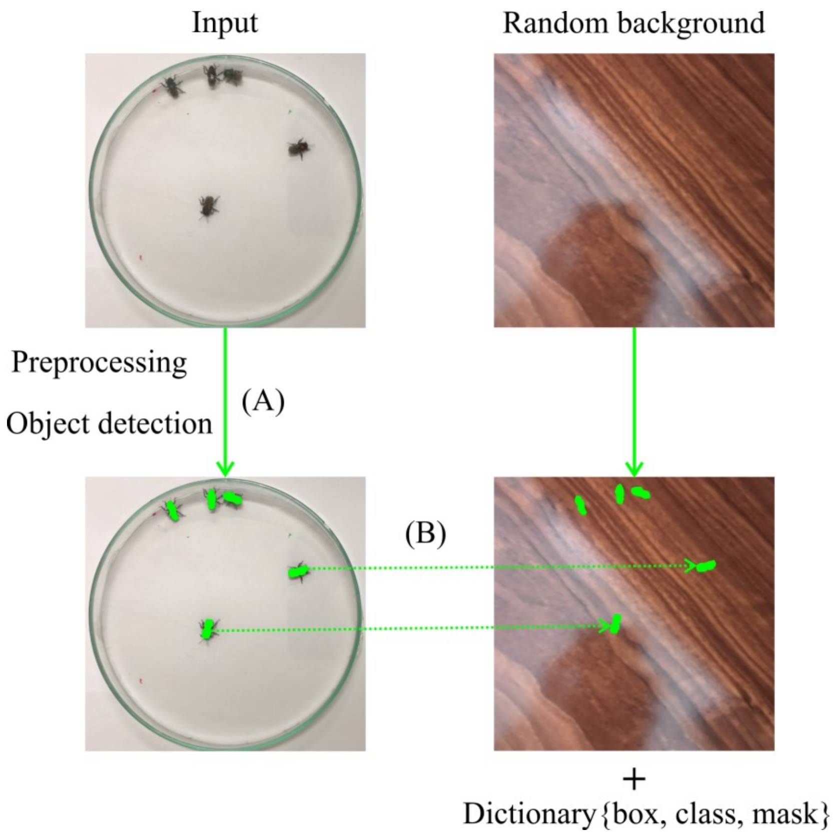


Figure S3. Schematic representation of the heuristic used to automatically generate labeled images for the IS model. The segmented objects (indicated by green masks) on a homogeneous background are glued to random backgrounds.

#### A.4 Post-processing

When the threshold is set manually or using Otsu's method, post-processing with different morphological operations is applied to eliminate residues. First, dilation is used to fill parts that belong to the same individual but are detected separately. Second, the gradient is calculated and subtracted from the expanded frame to eliminate undesirable edges. Finally, erosion is applied to eliminate any noise erroneously detected as individuals. When thresholding is performed by IS, morphological operations are not required, as the proposed AI model usually does not detect the undesirable parts of fragmented individuals.

#### A.5 Position and identity

The vectors containing the points of each contour of the objects are identified in this step. Contours are identified without establishing hierarchies, while retaining only the extreme points of the contour line segments. Contour measurements such as the area, length, and the ratio between the area and length are calculated to restrict the contours that are identified based on the user's inputs. Among the identified contours, the centroid and Cartesian position of each individual are determined.

When the number of contours identified is smaller than the number of individuals specified by the user, the nonhierarchical clustering k-means algorithm is applied to separate merged individuals. The number of groups (*k*) in which the set of pixels will be grouped is equal to the number of individuals specified by the user. The initial *k* centroids are randomly defined among the set of data points. Then, the next set of centroids are chosen according to the probability of spreading between the centers (Arthur & Vassilvitskii, 2006). The contour points are compared with each centroid and are allocated to the group whose Euclidean distance is minimal. Considering the inputs for the algorithm $X = \{x_1, \ldots, x_n\}$ of *n* data points, this algorithm runs interactively to find a set $C = \{c_1, \ldots, c_k\}$ that minimizes the function $\varphi_x(C)$ as follows:

$$\varphi_x(C) = \sum_{x \in X} d\,(x, C)^2, \tag{13}$$

where $d(x, C)^2$ is the distance from *x* to the closest center in *C*. To choose centroids in the k-means algorithm, the first set of centers $C_0$ are randomly selected from the dataset. Then, this step is repeated for $2 \leqslant i \leqslant k$: $c_i$ is chosen to be equal to a data point $x_n$ according to the probability (Arthur & Vassilvitskii, 2006):

$$\frac{d(x_0, C)^2}{\varphi_x(C)}. \tag{14}$$

A combinatorial optimization algorithm is applied to the centroids to maintain the identity of individuals. This is based on the Euclidean distance between the set of centroids of the objects in the $frame_{i+1} = \{a_1, a_2, \ldots, a_n\}$ and the set of centroids in the $frame_i = \{b_1, b_2, \ldots, b_n\}$. Considering that each $a_n$ is assigned to only one $b_n$, the goal is to minimize the total cost of assignments about the distance matrix (*D*) between each $a_n$ and $b_n$:

$$D = \begin{bmatrix} d_{1,1} d_{1,2} \cdots d_{1,n} \\ d_{2,1} d_{2,2} \cdots d_{2,n} \\ \vdots \quad \vdots \quad \vdots \\ d_{n,1} d_{n,2} \cdots d_{n,n} \end{bmatrix}.$$

The mathematical model (Kuhn, 1955) for the assignments is given as $Minimum$: $\sum_{x=1}^{n} \sum_{j=1}^{n} d_{ij}$, where $d_{ij}$ is the cost (Euclidean distance) from centroid $a_n$ to centroid $b_n$. There are *n!* ways to assign $a_n$ to $b_n$, to achieve the optimal assignment, interactively, with the following steps:

1. The minimum of each row is subtracted from the entire row.

2. The minimum of each column is subtracted from the entire column.

3. All zeros in the matrix are crossed with the minimum possible lines.

*If* crossing lines = $n$, then the optimal assignment is found.

*Else*:

To determine the smallest entry not crossed by any line,

Subtract this entry from each uncrossed row and add it to each crossed column.

Proceed to step 3.

#### A.6 Kinematic variables

Based on the position (coordinates $x$, $y$) of each individual in the video frames ($n$), Ethoflow computes various kinematic parameters associated with an individual and the group behavior. The distance an animal walks during tracking is defined as the tracked distance ($td$) (Eq. 15). Dividing $td$ by the total time of the video, the mean speed can be calculated. Ethoflow also calculates the maximum speed achieved by the animal.

$$td = \sum_{i=1}^{n} \sqrt{(x_{i+1} - x_i)^2 + (y_{i+1} - y_i)^2} \tag{15}$$

The turning angle ($ta$) is the absolute sum of the angles (∘) of the movement divided by the video frames ($n$) (Eq. 16), while the meandering ($M$) is divided by $td$ (Eq. 17). The angle of the movement is the arctangent of the locomotion in planes $y$ ($\Delta y_i$) and $x$ ($\Delta x_i$).

$$ta = \frac{1}{n} \sum_{i=1}^{n} \left| \left( \frac{arctan\left(\frac{\Delta y_i}{\Delta x_i}\right) 180}{\pi} \right) \right| \tag{16}$$

$$M = \frac{1}{td} \sum_{i=1}^{n} \left| \left( \frac{arctan\left(\frac{\Delta y_i}{\Delta x_i}\right) 180}{\pi} \right) \right| \tag{17}$$

The movement of individuals is categorized based on the user-defined values. When defining the analysis protocol, the user defines the thresholds for low ($tl$) and high movement ($th$). Thus, considering the movement of individuals in each frame as $mf$: $mf \leq tl$ is counted as resting; $tl < mf \leq th$ is counted as mean movement; $mf > th$ is counted as fast movement. The sum of these counts is divided by the frames per second (*fps*) used to sample the video to obtain these values in time.

The user also sets a threshold for interaction (*ti*). The interaction is considered when the individuals approach a distance $\leq ti$. The sum of all interactions of an individual is defined as the centrality. The network density (*nd*) is a measurement associated with group interaction (Eq. 18). A network is a set of items in which the vertices are defined as nodes (*n*), and the connections among them are defined as edges (*m*) (Newman, 2003). Here, the nodes are the individuals and the edges represent the number of interactions among them.

$$nd = \frac{2m}{n(n-1)} \quad (18)$$

If the user defines a region of interest (*ri*), Ethoflow computes how long the individuals stayed inside this region, considering the position (coordinates *x*, *y*) of each individual in the video frames (*n*):

$$\sum_{i=1}^{n} (x_i, y_i) \in ri. \quad (19)$$

Considering the direction unit (*u*) of the individuals (*i*), the proportion of the group polarized (*p*) at each frame is calculated as

$$p = \frac{1}{i}\left|\sum_{j=1}^{i} u_j\right|. \quad (20)$$

The angular momentum (rotate; *r*) for each frame is a cross product between the distance (*d*) of an individual to the center of mass of the group and the direction of movement (*u*):

$$r = \frac{1}{i}\left|\sum_{j=1}^{i} u_j \times r_j\right|. \quad (21)$$

These parameters provide information on the global structure of the group (Tunstrøm et al., 2013), such as the polarization (*gp*), swarming (*gs*), milling (*gm*), and transition (*gt*). The sum of these counts is divided by the *fps* to obtain these values in time:

$$gp = \frac{\sum p > 0.65 \text{ and } r < 0.35}{fps}, \quad (22)$$

$$gs = \frac{\sum p < 0.35 \text{ and } r < 0.35}{fps}, \quad (23)$$

$$gm = \frac{\sum p < 0.35 \text{ and } r > 0.65}{fps}, \tag{24}$$

$$gr \neq \{gp, gs, gm\}. \tag{25}$$

#### A.7 Specific behavior

Different hyperparameters were tested to find a suitable convolutional neural network (CNN) model. This model was trained to recognize trophallaxis in the stingless bees *Melipona quadrifasciata*, and can be used to recognize many binary behaviors. Using the validation accuracy of such a variable response, the interaction between dropout and the activation function of the network output was evaluated (Fig. S4A). Dropout is a regularization technique that randomly zeros out the input units of a layer, breaking fixed patterns to avoid overfitting (Srivastava, Hinton, Krizhevsky, Sutskever, & Salakhutdinov, 2014). Here, a better response was obtained with an intermediate dropout value of 0.2 (Fig. S4A). The higher rates, despite reducing overfitting, decreased the accuracy. The activation function of the network output that presented the best response was the sigmoid function (Eq. 26). This function binarizes the network output (0 or 1). As it involves a binary behavior classification, the sigmoid function is expected to generate a better output.

$$\left\{ f(x) = \frac{1}{1+e^{-x}}, \text{where } x \text{ is the output from the previously hidden layer} \right\}. \tag{26}$$

The activation function of the inner layers and the optimization method were also evaluated. The best result was obtained with the exponential linear unit (Elu) function, along with mini-batch stochastic gradient descent (mini-batch SGD) as the optimizer (Fig. S4B). The outputs of the layers pass through an activation function. These functions introduce nonlinearity into the network, which increases the dimensionality of the representation space and accords meaning to the stacking in the sequence of network layers. The Elu function (Eq. 27) is an identity function for positive values, and it tends smoothly to $-\alpha$ for negative values. This function saturates for very small (extremely negative) values, resulting in the activation average being close to zero. Thus, ELUs tend to normalize the output of the layer, accelerate learning, and increase the accuracy (Clevert, Unterthiner, & Hochreiter, 2015).

$$f(x) = \left( \begin{array}{c} x, \ \ if\ x > 0 \\ \alpha * exp(x) - 1, \ \ if\ x \leqslant 0 \end{array} \right). \tag{27}$$

In the mini-batch SGD, the term stochastic refers to a random sampling of lots in the data. Based on the loss value, the optimizer plays the role of updating the trainable parameters of the network (weights). This is executed by calculating the loss gradient in relation to the parameters (current weights) of the network. Mathematically, this process is performed by deriving the cost function and finding the gradient for the current weights. Then, the weights are updated in the opposite direction of the gradient, reducing the loss slightly with each batch. Because the classification is binary (the output from the network is a probability), binary cross-entropy (Eq. 28) was used as the cost function. Cross-entropy is a measure of the distance between the expected result $y$ and the predictions $p(y)$.

$$H_p(q) = \frac{-1}{n}\sum_{i=1}^{n} y_i * log\big(p(y_i)\big) + (1 - y_i) * log\big(1 - p(y_i)\big), \qquad (28)$$

where $n$ is the number of network outputs.

As an increase in the learning rate tended to decrease the accuracy (Fig. S4C), the lowest rate tested (0.0005) was maintained. The learning rate determines the magnitude of gradient descent. At high learning rates, network updates can result in great randomness.

The network interacts with the data in mini-batches, i.e., it does not process an entire dataset simultaneously; rather, the data is divided into small batches. Although this hyperparameter is important in CNN models (Radiuk, 2017), it does not play an important role in our model (Fig. S4D). Therefore, one of the smallest values (batch size = 5) was selected to accelerate the training time of the network.

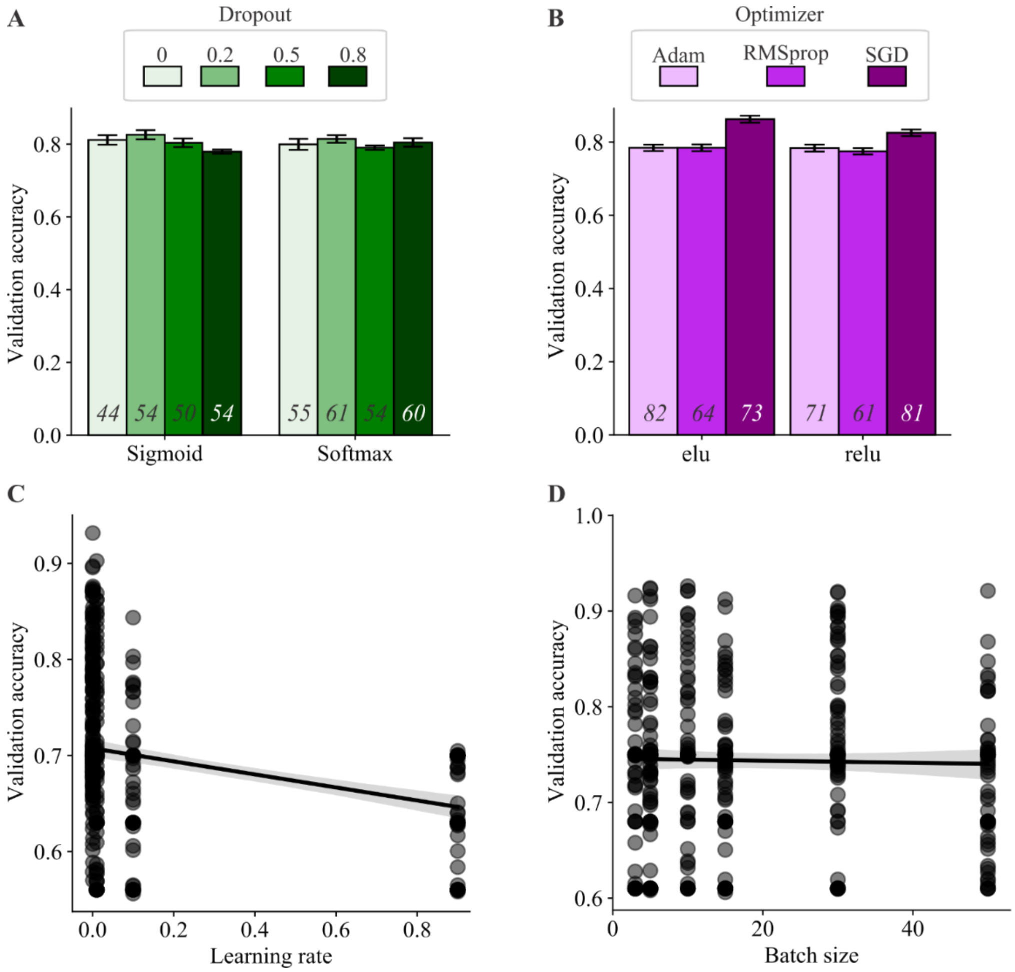


Figure S4: Hyperparameter optimization of the CNN model used in Ethoflow. (A) Compilation of the validation accuracy in response to dropout and the activation function of the network output. (B) Validation accuracy in response to the optimizer and activation function of the inner layers. (A), (B) The bars represent the mean ± standard error. The values at the base of each bar represent the number of times a given configuration was tested. Scatterplot of the accuracy as a function of the (C) learning rate and (D) batch size. The translucent band around the line of regression represents the confidence interval ($n$ = 432).

In many statistical models, the normalization of variables is important (for e.g., to avoid the predominance of some variables due to different scales). To this end, batch normalization layers were used in the CNN model. This layer can adaptively normalize the data as the mean and variance change during training (Ioffe & Szegedy, 2015).

Using the parameters defined above, the size of the network (number of layers) was also evaluated, and better accuracy was obtained with smaller architectures (Table *1*). While the presence of more layers (a higher-dimensional representation space) allows the network to learn more complex representations, this increases the computational cost of the network; accordingly, model L7 was employed.

Table 1: Different architectures tested to ascertain the ideal number of layers in the CNN model ($n = 28$).

| | Number of layers | | |
|---|---|---|---|
| **Validation accuracy (mean ± sd)** | **Convolutional** | **Dense** | **Model** |
| 0.63 ± 0.028 | 5 | 4 | L1 |
| 78 ± 0.036 | 5 | 3 | L2 |
| 0.83 ± 0.042 | 5 | 2 | L3 |
| 0.81 ± 0.063 | 4 | 4 | L4 |
| 0.8 ± 0.121 | 4 | 2 | L5 |
| 0.8 ± 0.020 | 3 | 4 | L6 |
| 0.91 ± 0.031 | 3 | 3 | L7 |

Data augmentation is a powerful technique for mitigating overfitting. Using the defined architecture (model L7 in Table *1*), different configurations of data augmentation were tested (Table *2*). Excessive data augmentation reduces the accuracy, while sets with little augmentation increase overfitting. Thus, set 3 was deemed the best option to address the problem of overfitting.

Table 2: Sets tested for data augmentation. In all the tests, the horizontal flip and fill mode = "nearest" was used. Model L7 in table A.1 was used for these tests.

| Parameters | Set 1 | Set 2 | Set 3 | Set 4 |
|---|---|---|---|---|
| Rotation range | 20 | 16 | 14 | 11 |
| Width shift range | 0.1 | 0.08 | 0.06 | 0.01 |
| Height shift range | 0.1 | 0.08 | 0.06 | 0.01 |
| Shear range | 0.05 | 0.02 | 0.01 | 0.008 |
| Zoom range | 0.1 | 0.08 | 0.06 | 0.01 |

The annotated images used to train the CNN model for recognizing trophallaxis were automatically generated through a heuristic. When bees perform trophallaxis, they position themselves in front of each other and exchange food. Based on this predictable positioning, the heuristic was based on the area and body length of the individuals. Initially, the program estimates the median ($M$) and standard deviation (*sd*) of the body area (*a*) and length (*l*) in frames where there is no crossing (no meeting between individuals). Subsequently, the software obtains the images (*b*) from the video and labels them as trophallaxis *if*:

$$\left\{\begin{matrix} area(b) \geqslant 2 * \big(M(a) - sd(a)\big) \text{ and} \\ area(b) \leqslant 2 * \big(M(a) + sd(a)\big) \text{ and} \\ length(b) \geqslant 2 * \big(M(l) - sd(l)\big) \text{ and} \\ length(b) \leqslant 2 * \big(M(l) + sd(l)\big) \end{matrix}\right\}, Else\text{: } b \text{ is not trophallaxis.}$$

#### A.8 Output

The behavioral parameters are automatically saved in a comma-separated values (csv) file in the path defined by the user. This file also contains the raw data, which are the coordinates (x, y) of movement. Using this data, the user is free to calculate other kinematic parameters, in addition to those automatically computed by the software.

The accuracy of the assessments is measured using the detection rate (*dr*). Given the instantaneous speed vector $IS = (is_1, \dots, is_n)$, where $n$ is the number of frames in the video, *dr* is defined as:

$$dr = 1 - \left(\left(\frac{\sum_{i=1}^{n} is_i > 2 * P_{.95}(IS)}{is_i}\right)\left(\frac{1}{n}\right)\right), \quad (29)$$

where $P_{.95}$ is the percentile at 95% of the *IS* vector.

### B. Performance and applications (section B)

#### B.1 Artificial intelligence models

The Ethoflow software was run on a Dell G3 machine, with Linux Ubuntu 19.4 64 bits, Intel i7-9750H CPU 2.60 GHz x 12, 8GB RAM and GPU NVIDIA® GeForce® GTX 1660 (6GB) Ti Max-Q. Applying the heuristic to the IS model (Section A.3), 1,325 labeled images were generated to train the model for tracking the stingless bee *M. quadrifasciata* (Fig. S5). Of these data, 976 (74%) images were used for training, 249 (19%) for validation, and 100 (7%) to evaluate the classification using the AP. To train the CNN model for the recognition of trophallaxis in *M. quadrifasciata*, 1,270 labeled images were generated (724 for non-trophallaxis and 546 for trophallaxis) (Fig. S6). In this dataset, 70% of the data was used for training, while 20% was used for validation. In addition, another sample dataset (10%) was used to assess the classifier's performance based on the global accuracy from the confusion matrix, Kappa index, and Z-test (5%).

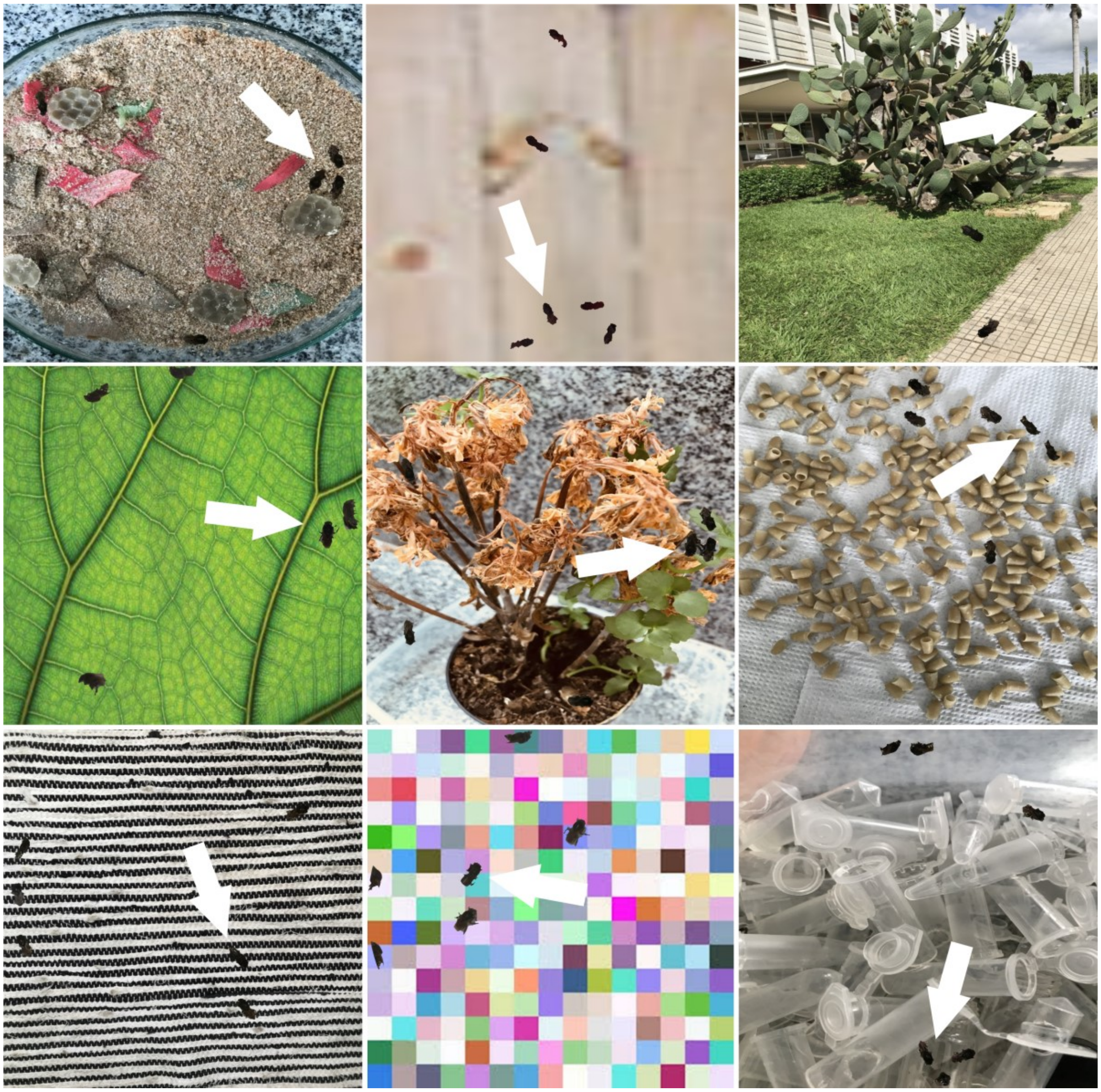

Figure S5. Examples of some images that were generated automatically to train the model for tracking the stingless bee *Melipona quadrifasciata* in different conditions of a heterogeneous background. The white arrows indicate some bee contours pasted in the backgrounds.

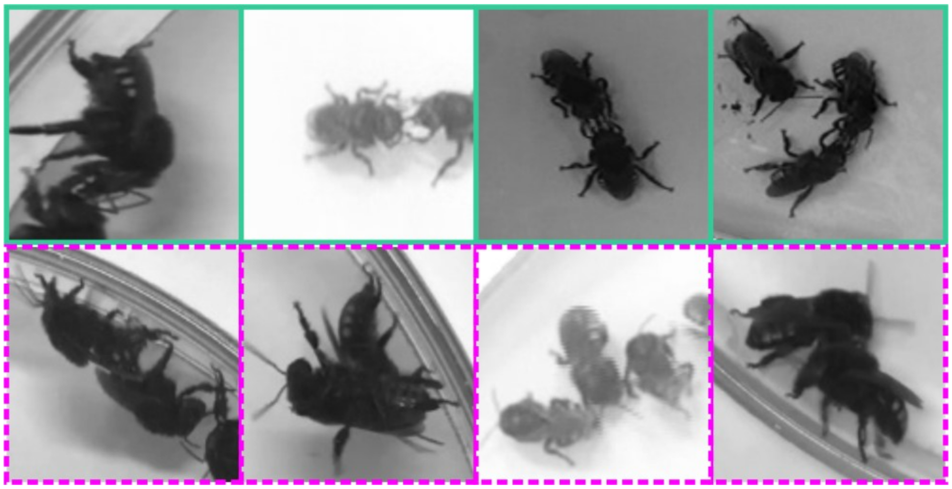

Figure S6. Examples of images that were automatically labeled with our heuristic to train the CNN model for the recognition of trophallaxis in the stingless bee *Melipona quadrifasciata*. The images with green outlines (top panel) are examples of trophallaxis. The images with dashed purple outlines (bottom panel) are examples of non-trophallaxis.

#### B.2 Application of the proposed software in behavioral assessments

To conduct behavioral assessments using Ethoflow, bees of both species were collected from four colonies each of *M. quadrifasciata* and *Partamona helleri* in Viçosa county, State of Minas Gerais, Brazil (20° 45′ S and 42° 52′ W). The collected bees were kept for 1 h in the laboratory under conditions similar to those found in their colonies (28 ºC and 80% relative humidity in total darkness) (Botina et al., 2020). Subsequently, bee behavior was recorded in the arenas (Petri dish, 9 cm diameter, 2 cm height) for 15 min with a digital video camera (HDR-XR520V, Sony Corporation) at 30 fps and high definition (1920 × 1080 pixels). Behavioral bioassays were performed in a room with artificial fluorescent light at 25 ± 3 °C and 70 ± 5% relative humidity. Bioassays were performed with 37 replicates, with each replicate corresponding to a group of five bees of the two species. The kinematic variables measured with Ethoflow included centrality, polarization, milling, resting, meandering, and tracked distance. In the centrality response, the interaction was considered when the individuals approached a distance ≤ 1.41 cm. An instantaneous tracked distance ≤ 0.046 cm $frame^{-1}$ was counted as resting. Centrality was the response variable in the model with interaction between polarization and bee species, or model with interaction between milling and bee species. Meandering was the response variable in the model with interaction between resting and bee species. In addition, the tracked distance between the bee species was compared. These models were fitted with generalized linear models (GLM) with a gamma distribution, displaying adequate distribution for continuous data in which the variance increases with the square of the mean (Crawley, 2012). When an explanatory variable had no significant effect, the model was simplified, and the results were plotted as a function of the significant variable.

In the toxicological bioassay employed to assess changes in trophallaxis behavior, *M. quadrifasciata* bees were collected and acclimated as previously described. Then, these bees were orally exposed to the commercial formulation (cf) (water-dispersible granules at 700 g a.i. $Kg^{-1}$, Bayer CropScience, São Paulo, SP, Brazil) of the neonicotinoid imidacloprid in a sublethal concentration (0.2 mg cf $L^{-1}$). This concentration is 300× smaller than that recommended for the control of the whitefly *Bemisia tabaci* (60 mg cf $L^{-1}$) (MAPA, 2020). The pesticide imidacloprid is commonly associated with bee decline and causes motor impairments in bees (Lima, Martins, Oliveira, & Guedes, 2016). After 3 h of exposure, the bees were filmed as previously described, and trophallaxis behavior was quantified using Ethoflow. Trophallaxis response (n = 60) to the pesticide was assessed using a GLM with a Poisson distribution, which is a suitable distribution for count data (Crawley, 2012).

#### B.3 Performance of the developed software

Some software features were evaluated during behavioral assessments (Section B.2). In addition, other videos with varying resolution, number of individuals, animals, and backgrounds were also evaluated (Fig. S7). To assess whether the fps responds to the interaction between the resolution and number of individuals, a multiple regression model was applied. The effect of centrality and the number of individuals in fps was also assessed using a GLM with a gamma distribution. Analysis of covariance (ANCOVA) was performed to assess whether the detection rate varied with the interaction between the number of individuals and background (homogeneous and heterogeneous).

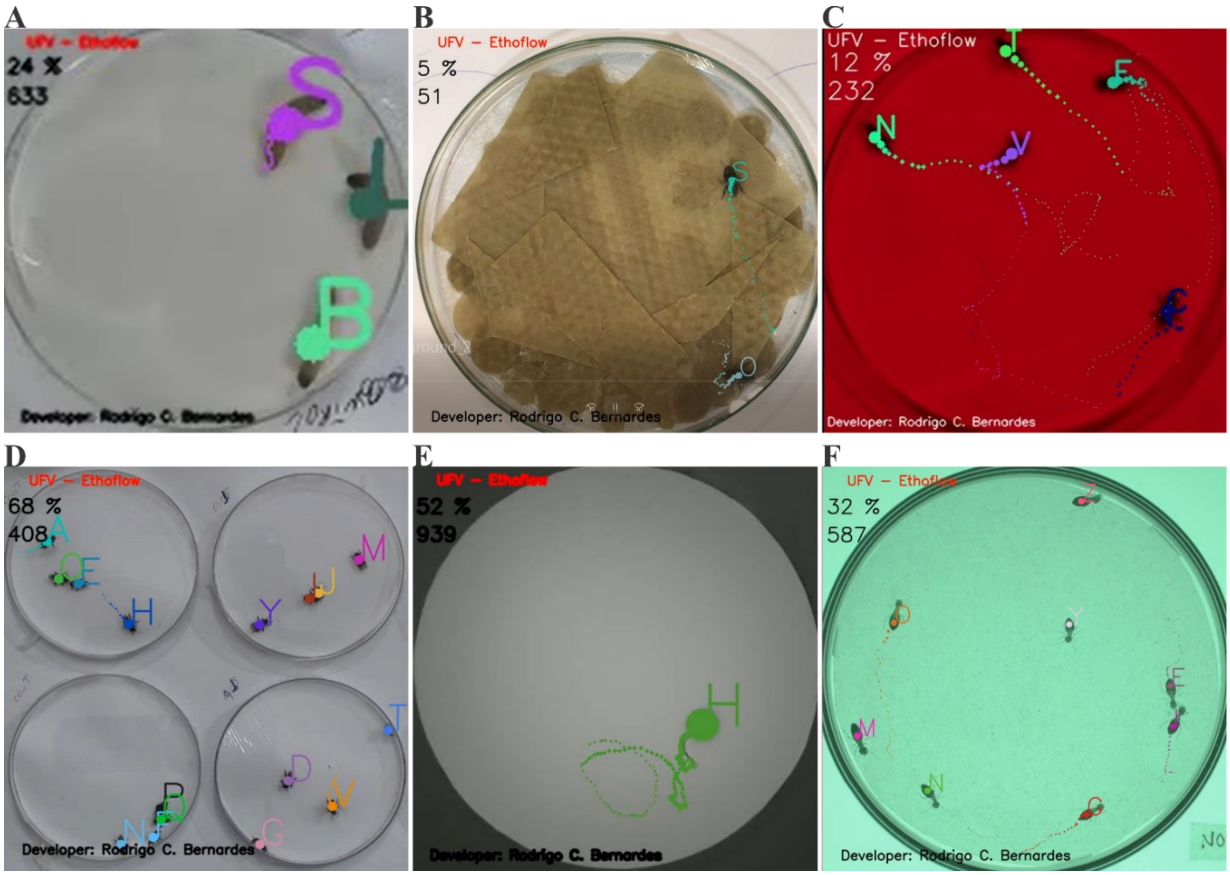


Figure S7. Examples of videos processed with Ethoflow. (A) Monitoring of *Galleria mellonella* larvae in the low-quality video. (B), (C) and (D) Monitoring bees (*Melipona quadrifasciata* and Partamona helleri) in different background and luminosity conditions. Monitoring of the C57Bl/6 mouse strain (E) and termites (*Constrictotermes cyphergaster*) (F). (E) and (F) were obtained from the freely available GitHub repository (https://github.com/vivekhsridhar/tracktor/tree/master/examples) (Sridhar, Roche, & Gingins, 2019).

#### B.4 Videos

Representative videos processed with Ethoflow have been submitted at https://github.com/bernardesrodrigoc/Ethoflow; DOI: https://doi.org/10.5281/zenodo.3956831

#### B.5 References

Abdulla, W. (2017). Mask R-CNN for object detection and instance segmentation on Keras and

TensorFlow. *GitHub Repository*. Github.

Arthur, D., & Vassilvitskii, S. (2006). *k-means++: The Advantages of Careful Seeding*. Stanford. Retrieved from http://ilpubs.stanford.edu:8090/778/

Botina, L. L., Bernardes, R. C., Barbosa, W. F., Lima, M. A. P., Guedes, R. N. C., & Martins, G. F. (2020). Toxicological assessments of agrochemical effects on stingless bees (Apidae, Meliponini). *MethodsX*, 100906. doi:https://doi.org/10.1016/j.mex.2020.100906

Clevert, D.-A., Unterthiner, T., & Hochreiter, S. (2015). Fast and Accurate Deep Network Learning by Exponential Linear Units (ELUs). Retrieved from http://arxiv.org/abs/1511.07289

Crawley, M. J. (2012). *The R book* (2nd ed.). Chichester: Wiley. Retrieved from https://www.wiley.com/en-us/The+R+Book%2C+2nd+Edition-p-9780470973929

Everingham, M., Van Gool, L., Williams, C. K. I., Winn, J., & Zisserman, A. (2010). The Pascal Visual Object Classes (VOC) Challenge. *International Journal of Computer Vision*, *88*(2), 303–338. doi:10.1007/s11263-009-0275-4

He, K., Gkioxari, G., Dollár, P., & Girshick, R. (2018). Mask R-CNN. *IEEE Transactions on Pattern Analysis and Machine Intelligence*, *42*(2), 386–397. doi:10.1109/TPAMI.2018.2844175

He, K., Zhang, X., Ren, S., & Sun, J. (2016). Deep residual learning for image recognition. In *Proceedings of the IEEE Computer Society Conference on Computer Vision and Pattern Recognition* (Vol. 2016-December, pp. 770–778). IEEE Computer Society. doi:10.1109/CVPR.2016.90

Ioffe, S., & Szegedy, C. (2015). Batch Normalization: Accelerating Deep Network Training by Reducing Internal Covariate Shift.

Kuhn, H. W. (1955). The Hungarian method for the assignment problem. *Naval Research Logistics Quarterly*, *2*(1–2), 83–97.

Lima, M. A. P., Martins, G. F., Oliveira, E. E., & Guedes, R. N. C. (2016). Agrochemical-induced stress in stingless bees: peculiarities, underlying basis, and challenges. *Journal of Comparative Physiology A: Neuroethology, Sensory, Neural, and Behavioral Physiology*, *202*(9–10), 733–747. doi:10.1007/s00359-016-1110-3

MAPA. (2020). Ministério da Agricultura, Pecuária e Abastecimento (MAPA). Retrieved 30 June 2020, from http://agrofit.agricultura.gov.br/agrofit_cons/principal_agrofit_cons

Newman, M. E. J. (2003). The Structure and Function of Complex Networks. *SIAM Review*, *45*(2), 167–256. doi:10.1137/S003614450342480

Otsu, N. (1979). A Threshold Selection Method from Gray-Level Histograms. *IEEE Transactions on Systems, Man, and Cybernetics*, *9*(1), 62–66. doi:10.1109/TSMC.1979.4310076

Radiuk, P. M. (2017). Impact of Training Set Batch Size on the Performance of Convolutional Neural Networks for Diverse Datasets. *Information Technology and Management Science*, *20*(1), 20–24. doi:https://doi.org/10.1515/itms-2017-0003

Sridhar, V. H., Roche, D. G., & Gingins, S. (2019). Tracktor: Image-based automated tracking of animal movement and behaviour. *Methods in Ecology and Evolution*, *10*(6), 815–820. doi:10.1111/2041-210X.13166

Srivastava, N., Hinton, G., Krizhevsky, A., Sutskever, I., & Salakhutdinov, R. (2014). Dropout: A Simple Way to Prevent Neural Networks from Overfitting. *Journal of Machine Learning Research*, *15*(56), 1929–1958. Retrieved from http://jmlr.org/papers/v15/srivastava14a.html

Tunstrøm, K., Katz, Y., Ioannou, C. C., Huepe, C., Lutz, M. J., & Couzin, I. D. (2013). Collective States, Multistability and Transitional Behavior in Schooling Fish. *PLoS Computational Biology*, *9*(2), e1002915. doi:10.1371/journal.pcbi.1002915
